# Genomic characterisation of a novel species of *Erysipelothrix* associated with mortalities among endangered seabirds

**DOI:** 10.1101/2022.06.23.497316

**Authors:** Jiadong Zhong, Matej Medvecky, Jérémy Tornos, Augustin Clessin, Hubert Gantelet, Amandine Gamble, Taya L. Forde, Thierry Boulinier

## Abstract

Infectious diseases threaten endangered species, particularly in small isolated populations. Seabird populations on the remote Amsterdam Island in the Indian Ocean have been in decline for the past three decades, with avian cholera caused by *Pasteurella multocida* proposed as the primary driver. However, *Erysipelothrix* spp. has also been sporadically detected from albatrosses on Amsterdam Island and may be contributing to some of the observed mortality. In this study, we genomically characterised 16 *Erysipelothrix* spp. isolates obtained from three Indian yellow-nosed albatross chick carcasses in 2019. Two isolates were sequenced using both Illumina short-read and MinION long-read approaches, which – following hybrid assembly – resulted in closed circular genomes. Mapping of Illumina reads from the remaining isolates to one of these new reference genomes revealed that all 16 isolates were closely related, with a maximum of 13 nucleotide differences distinguishing any pair of isolates. The nucleotide diversity of isolates obtained from the same or different carcasses was similar, suggesting all three chicks were likely infected from a common source. These genomes were compared with a global collection of genomes from *E. rhusiopathiae* and other species from the same genus. The isolates from albatrosses were phylogenetically distinct, sharing a most recent common ancestor with *E. rhusiopathiae*. Based on phylogenomic analysis and standard thresholds for average nucleotide identity and digital DNA-DNA hybridisation, these isolates represent a novel *Erysipelothrix* species, for which we propose the name *Erysipelothrix amsterdamensis* sp. nov. The type strain is *E. amsterdamensis* A18Y020d^T^. The implications of this bacterium for albatross conservation will require further study.

## Introduction

Infectious diseases are among the major threats to the conservation of endangered species, especially at local scales [1]. This is particularly true for populations in remote areas, where anthropogenic activity has historically been limited but is increasing, such as with commercial tourists to the Antarctic [2, 3]. Concerns have been raised about disease-associated declines among albatross populations and how this could influence their long-term sustainability [4]. Avian cholera, caused by the bacterium *Pasteurella multocida*, has been causing recurrent chick mortalities among two endangered albatross species on the remote Amsterdam Island in the southern Indian Ocean since at least the mid-1990s when it was first detected [5–7]. Both the Indian yellow-nosed albatross (*Thalassarche carteri*) and the sooty albatross (*Phoebetria fusca*) have been affected, and the endemic endangered Amsterdam albatross (*Diomedea amsterdamensis*) is also at risk [5]. The bacterium *Erysipelothrix rhusiopathiae* has also been detected by PCR in swabs from live birds of these three albatross species and from the endangered northern rockhopper penguin (*Eudyptes moseleyi*) on Amsterdam Island [6], and it has been isolated concomitantly from mortalities among the Indian yellow-nosed albatross [5]. During ongoing population health monitoring conducted to infer transmission dynamics and management opportunities [8–10], our team isolated *Erysipelothrix* spp. (identified at the time as *E. rhusiopathiae*) in 2019 from the carcasses of three Indian yellow-nosed albatross chicks on the island, strengthening the hypothesis that this bacterium could also be contributing to albatross mortalities and decreased breeding success.

*E. rhusiopathiae* is a small, Gram-positive bacillus. While best known as an opportunistic pathogen of pigs and poultry [11, 12], it affects a wide range of species [13] and in recent years it has been implicated in large-scale wildlife mortalities [14]. It was also responsible for jeopardising the translocation success of several critically endangered kakapo – a flightless parrot – between New Zealand and offshore islands [15], and has caused the death of other endangered birds in that area [16]. Erysipelas in wild birds is a septicaemic disease [12], and deaths have been reported in a range of wild terrestrial birds and seabirds worldwide [15]. Mortalities occur sporadically, and tend to be limited to a small number of individuals, although cases are likely under-recognised and under-reported [17]. Rare mass mortality events in wild birds have also been observed [18].

In this study, we sought to genomically characterise the *Erysipelothrix* strains isolated from Indian yellow-nosed albatross on Amsterdam Island to identify potential sources, and to assess the relationship between the multiple observed cases on the island.

## Methods

Between December 2018 and March 2019, 27 chick and 2 adult carcasses of Indian yellow-nosed albatross were necropsied in the field on Amsterdam Island (−37.797135, 77.571521) as part of long-term population health monitoring (Appendix S1, Figure S1, Table S1). Samples from multiple organs (liver, lung, brain, and heart) were collected from the carcasses for bacteriological and histological analysis. Isolates of *Erysipelothrix* spp. were identified from three of the chick carcasses, all sampled between the 13^th^ and the 16^th^ of January, 2019. The bacterial growth in the field (Appendix S1) showed *Erysipelothrix* spp. exclusively, suggesting that this bacterium could have been the cause of death. Between four and six isolates were obtained from each carcass (n = 16 total). These were initially identified in the laboratory as *E. rhusiopathiae* based on colony morphology on horse blood agar and MALDI-TOF, with the exception of strain A18Y019b, which was classified as *Erysipelothrix* spp. DNA was extracted from sub-cultured isolates (loop of colonies) using the GenElute Bacterial Genomic DNA Kit (Sigma-Aldrich) as per the manufacturer’s instructions. Extracted DNA was tested with a probe-based qPCR targeting *E. rhusiopathiae* [19], then sequenced at MicrobesNG (Birmingham, UK). Libraries were prepared using the Nextera XT v2 kit (Illumina) and sequenced on the Illumina HiSeq platform, generating 250 base pair paired-end reads. Long-read MinION sequencing (Oxford Nanopore) was performed at the University of Glasgow for two isolates (A18Y016a and A18Y020d), with library preparation done using the rapid barcoding kit (SQK-RBK004). Illumina reads were adapter and quality trimmed using Trimmomatic [20] and assembled *de novo* using SPAdes [21] by MicrobesNG. Assembled genomes were evaluated using the online version of Quality Assessment Tool for Genome Assemblies (QUAST) (http://cab.cc.spbu.ru/quast/) [22]. Unless otherwise indicated, the remaining bioinformatics analyses were performed within the CLIMB computing platform for microbial genomics [23]. To generate circularised (closed) reference genomes, the Illumina and MinION sequence data were jointly assembled by Unicycler v0.4.4 [24] applying the ‘normal’ (default) hybrid mode. For isolate A18Y016a, this did not result in a closed genome, so subsequently Canu v1.7 [25] was used to perform an initial assembly using the long-read MinION data, and Pilon v1.22 [26] applied to polish the assembly using Illumina reads, including three iterations. Tablet v1.21.02.08 [27] was used to visually check the distance between forward and reverse reads across the start and end positions of the linearised chromosome to confirm the genomes’ circular nature.

Prokka v1.13 [28] and BLAST2GO [29] were used to predict coding sequences (CDS), tRNA, rRNA and tmRNA. The general feature format (.gff) files obtained by the two programmes were merged to build a complete .gff annotation file. The PHASTER server (https://phaster.ca/) [30] was used to identify and annotate prophage sequences. plasmidFinder server v2.1 (https://cge.cbs.dtu.dk/services/PlasmidFinder/) [31] was used to search for plasmid sequence data in the genomes against the replicon database.

BWA-MEM [32] with default settings was used to map Illumina reads of each albatross isolate against the A18Y020d closed genome, and the mpileup command in SAMtools v1.13 [33] was used to generate summary text files of coverages across the reference genome. The pileup2snp command in Varscan v2.4.4 [34] was used to generate a list of high-quality single nucleotide polymorphisms (SNPs) that were filtered by applying parameters ‘--min-coverage’ of 10, ‘--min-var-freq’ of 0.8 and ‘--min-avg-qual’ of 20. A custom R script was developed to generate a multi-fasta alignment file based on concatenated SNPs. Roary v3.12.0 [35], with default parameters setting, was used to construct a core gene alignment for a wider phylogenetic analysis, i.e. to place the albatross isolates within the context of global *Erysipelothrix* spp. genomes. This included 106 genomes: a) the 16 isolates from this study; b) 86 previously sequenced *E. rhusiopathiae* isolates, representing a diversity of geographic and host origins and spanning the clades described for this species (Clade 1 (n = 7), Clade 2 (n = 14), intermediate clade (n = 8), Clade 3 (n = 57)) [36]; c) two genomes of *Erysipelothrix piscisicarius* (formerly referred to as *E*. sp. strain 2) [37, 38]: the type strain 15TAL0474^T^ (GCA_003931795.1) [39], and EsS2-7-Brazil (GCA_016617655.1). *E. piscisicarius* is the most closely-related species to *E. rhusiopathiae* currently known; d) a recently deposited *Erysipelothrix* genome, reported as a new species, *Erysipelothrix* sp. Poltava (GCA_023221615.1); and e) *Erysipelothrix tonsillarum* strain DSM14972^T^ (GCA_000373785.1), which was used as an outgroup, since this was previously reported to be ancestral to these other species [36]. The SNP alignment and core gene alignment files were used to estimate phylogenetic trees for the 16 albatross isolates and the full set of *Erysipelothrix* spp. genomes, respectively, using RAxML v8.2.4 [40] with Maximum-Likelihood (ML) method implementing a GTR-GAMMA nucleotide substitution model, selected after using MrModeltest v2.4 [41].

To determine the relatedness of the albatross isolates from this study with other less well-described *Erysipelothrix* species available in the Genome Taxonomy Database (https://gtdb.ecogenomic.org/), a separate phylogenetic tree based on the *rpoB* gene (beta subunit of RNA polymerase) was estimated. This gene has been found to be a more suitable marker for *Erysipelothrix* species delineation than the widely used 16S rRNA gene [37]. Nucleotide sequences of *rpoB* genes were extracted from whole-genome assemblies of representative samples (one per species, n = 16 total), and aligned using L-INS-i method as implemented in MAFFT v7.313 [42]. Tree topology was then inferred by RAxML v8.2.4, implementing the GTR model of nucleotide substitution; robustness of the tree was assessed by performing 1000 bootstrap replicates. For completeness, the 16S rRNA gene sequence of the newly sequenced strain A18Y020d was extracted through blastn alignment and compared with 16S gene sequences from *E. rhusiopathiae* strain ATCC 19414^T^ (NR_040837.1), *E. piscisicarius* strain 15TAL0474^T^ (NR_170392.1), and *E. tonsillarum* strain DSM 14972^T^ (NR_040871.1). Two different genome relatedness indices were calculated in order to compare the *Erysipelothrix* spp. isolated from albatrosses with previously characterised *Erysipelothrix* species and determine the species assignment. Pairwise nucleotide-level comparisons were made using average nucleotide identity (ANI) Calculator for OrthoANIu [43] (https://www.ezbiocloud.net/tools/ani), and digital DNA-DNA hybridisation (dDDH) was performed using GGDC [44, 45] (https://ggdc.dsmz.de), using Formula 2. This formula is independent of genome length and is thus robust to the use of incomplete draft genome assemblies, as well as differences in gene content.

The output of Roary was used in Scoary v1.6.16 [46] to conduct a pan-GWAS (genome-wide association study) in order to identify any genes that might be unique to the albatross isolates, where the binary trait considered was ‘albatross origin’ yes/no. NCBI blastn searches were conducted on the Scoary results (i.e. genes identified as unique to the albatross isolates that were absent in other *Erysipelothrix* spp.) to verify their absence in other bacterial species. Primer sets typically used to amplify *Erysipelothrix* spp. [47] and *E. rhusiopathiae* specifically [19] were evaluated *in silico* against the complete assemblies using Geneious v11.0.5 [48] to check for any mismatches. Finally, to further characterise the *Erysipelothrix* spp. isolates from albatross, blastn searches of the *de novo* Illumina assemblies were conducted to determine *in silico* the serotype [49] and whether any *spa* genes (A, B or C) [50] were present, using methods previously described [51].

## Results

Sixteen isolates identified as *Erysipelothrix* spp. were recovered from the carcasses of three albatross chicks. Fifteen of these were initially identified as *E. rhusiopathiae* based on colony morphology and MALDI-TOF with 99.9% confidence, and the other strain was identified as an *Erysipelothrix* sp. Phenotypic characteristics determined during strain isolation are given in the species description below. All 16 isolates were genome sequenced on the Illumina platform, which yielded a high average depth of coverage of at least 58X for all genomes (Table S1). *De novo* genome assemblies comprised between 52 and 58 contigs per genome, with a total assembly length ranging from 1.80 to 1.83 megabases. Complete circular genomes were obtained for isolates A18Y020d^T^ (Figure S2; assembly GCA_940143175) – designated the type strain – and A18Y016a (assembly GCA_940143155), with lengths of 1,908,712 bp and 1,910,750 bp, respectively. This is approximately 120 kilobases (kb) longer than the Fujisawa *E. rhusiopathiae* reference genome, corresponding to ∼100 additional CDS (Table S2). Like the Fujisawa *E. rhusiopathiae* genome, both genomes from albatrosses had a GC content of 36.6%. Preliminary annotation of the two albatross-derived genomes using Prokka showed that this method predicted a large number of hypothetical proteins, amounting to approximately one-third of predicted CDS. Blast2GO facilitated the annotation of just over 1500 further CDS in each genome. Using PHASTER, an intact prophage region comprising 48 genes was detected in both genomes (Figure S3, Table S2). Based on global alignment using MUSCLE, this showed 43.6% nucleotide identity with the incomplete phage sequence in the *E. rhusiopathiae* Fujisawa reference genome (gene loci ERH_0581 to ERH_0629). No plasmids were detected in either genome.

Pairwise identity of the 16S rRNA gene sequence from strain A18Y020d^T^ with that of *E. rhusiopathiae* strain ATCC 19414^T^ was 99.9%; only two SNP differences were present across 1479 nucleotides (at positions 472 and 473; TC instead of CT). Only three further nucleotide differences distinguished these sequences from the 16S sequence of *E. tonsillarum* strain DSM 14972^T^ (99.8% pairwise identity). Eight gaps were found in the alignment of the 16S sequence from *E. piscisicarius* strain 15TAL0474^T^ compared with the others, which also shared one of the SNPs from A18Y020d^T^ at position 472, and had one additional nucleotide difference. The phylogenetic tree (Figure 1), based on 427 conserved core genes, showed that the 16 *Erysipelothrix* spp. isolates from albatrosses form a monophyletic clade distinct from previously characterised *Erysipelothrix* spp. genomes, which shares its most recent common ancestor with *E. rhusiopathiae*. Based on the *rpoB* gene phylogeny, other *Erysipelothrix* species discovered to date are more phylogenetically distant (Figure S4), with the exception of the recently deposited sequence for *Erysipelothrix* sp. Poltava; based on *rpoB* analysis, ANI/dDDH (Figure S5) and phylogenomic analyses (Figure S6), strain Poltava is concluded to belong to *E. rhusiopathiae* Clade 2.

**Figure 1.**
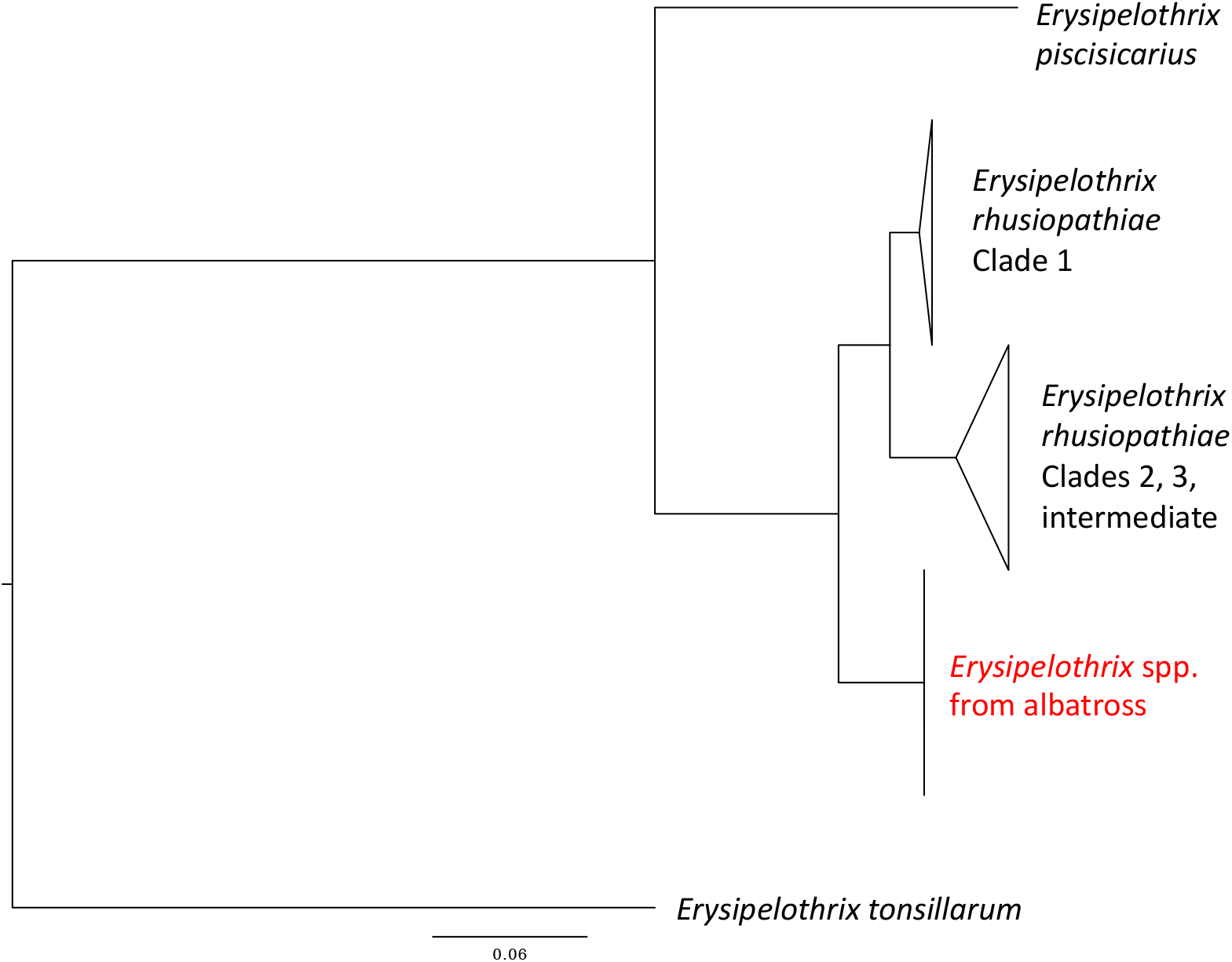
Maximum likelihood phylogenetic tree of *Erysipelothrix* spp. Isolates (n = 16) from Indian yellow-nosed albatross chicks on Amsterdam Island (red text) were compared with other representative *Erysipelothrix* genomes. The tree was estimated in RAxML using a multiple sequence alignment of conserved (core) genes identified using Roary, implementing a GTR-GAMMA model, and visualised using FigTree (http://tree.bio.ed.ac.uk/software/figtree/). *E. rhusiopathiae* clades are shown collapsed. Bootstrap values for all major nodes (those shown) were 100%. *E. tonsillarum* strain DSM14972^T^ was used as an outgroup for rooting.

Few SNP differences were observed among the albatross isolates. A total of 53 high-quality unique SNPs were detected across all 16 genomes when mapping against the closed reference genome (A18Y020d^T^; Figure 2). All genomes were unique, including where multiple isolates were sequenced from the same carcass (Table S1). The maximum pairwise SNP difference separating two isolates was 13. *In silico* serotyping based on a genomic polysaccharide biosynthetic locus [49] determined these isolates belong to serotype 1b. All genomes had identical sequences of the *spaA* gene, belonging to Group 2 [51]. These all had one unique amino acid residue at position 288 in comparison with any previously described sequences (serine instead of alanine).

**Figure 2.**
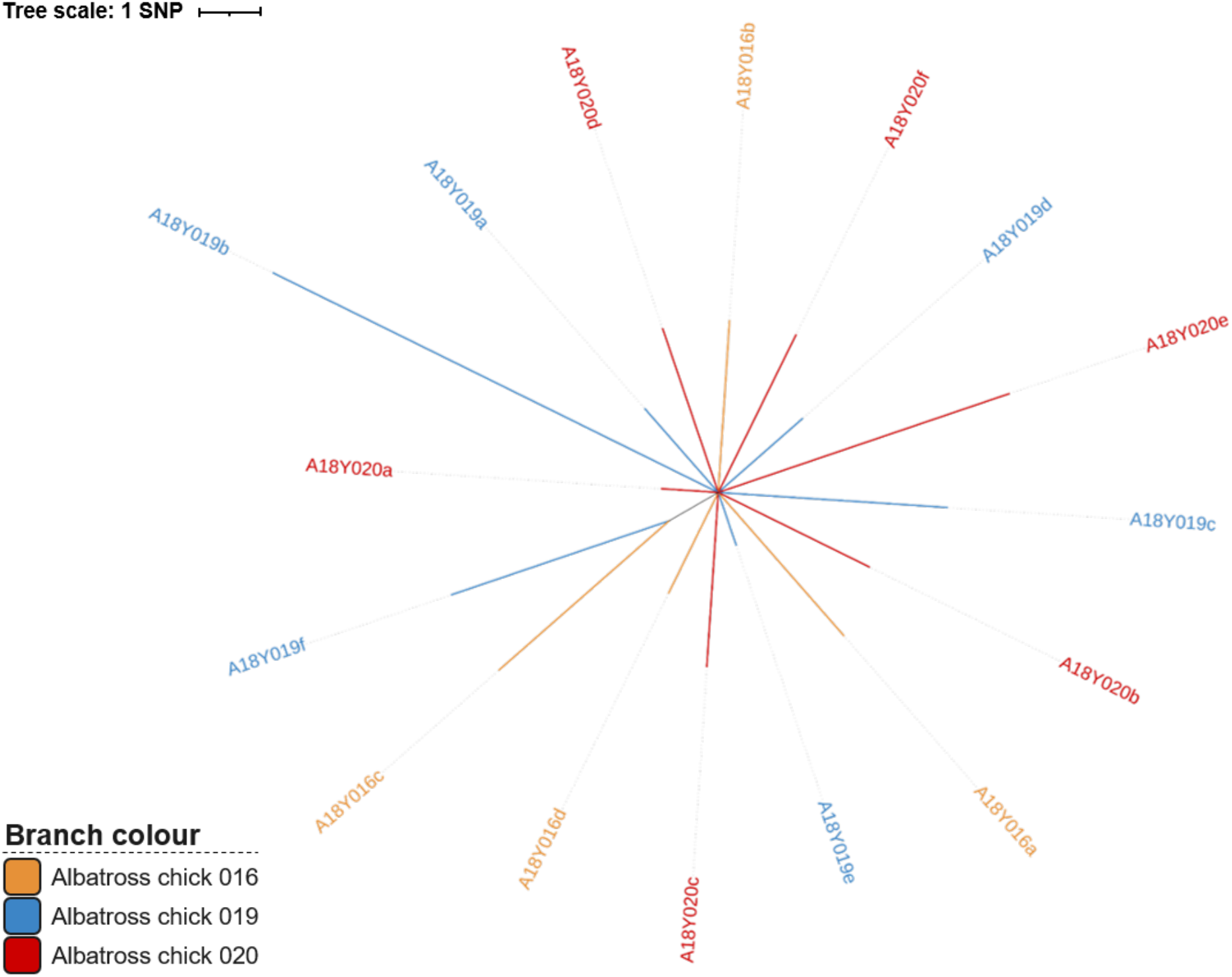
Unrooted maximum likelihood tree of 16 *Erysipelothrix* spp. isolates from Indian yellow-nosed albatross from Amsterdam Island. The tree was based on high-quality concatenated SNPs detected through mapping to the circularised genome of A18Y020d^T^. Isolates originated from three different carcasses, which are distinguished by different branch colours (as shown in the legend). Tree was plotted with Interactive Tree Of Life (iTOL) version 6.3.2 [52].

The ANI scores comparing the representative *Erysipelothrix* spp. type strain from albatross against *E. rhusiopathiae* were 95.0% with *E. rhusiopathiae* Clade 1, and just above 94% for representatives from the other clades, whereas they were 86.9% with *E. piscisicarius* and 80.9% with *E. tonsillarum* (Figure 3). The dDDH scores with *E. rhusiopathiae* Clades 1, 2 and 3 were 60.0%, 56.3% and 55.5%, respectively, while they were 32.7% with *E. piscisicarius* and 22.8% with *E. tonsillarum*. The accepted values for categorising isolates as the same species are >95% similarity for ANI [53, 54] and >70% for dDDH [45]. Thus, based on these standard metrics for genomic comparison, and the phylogenomic analysis, we conclude that these isolates belong to a novel species of the genus *Erysipelothrix*.

**Figure 3.**
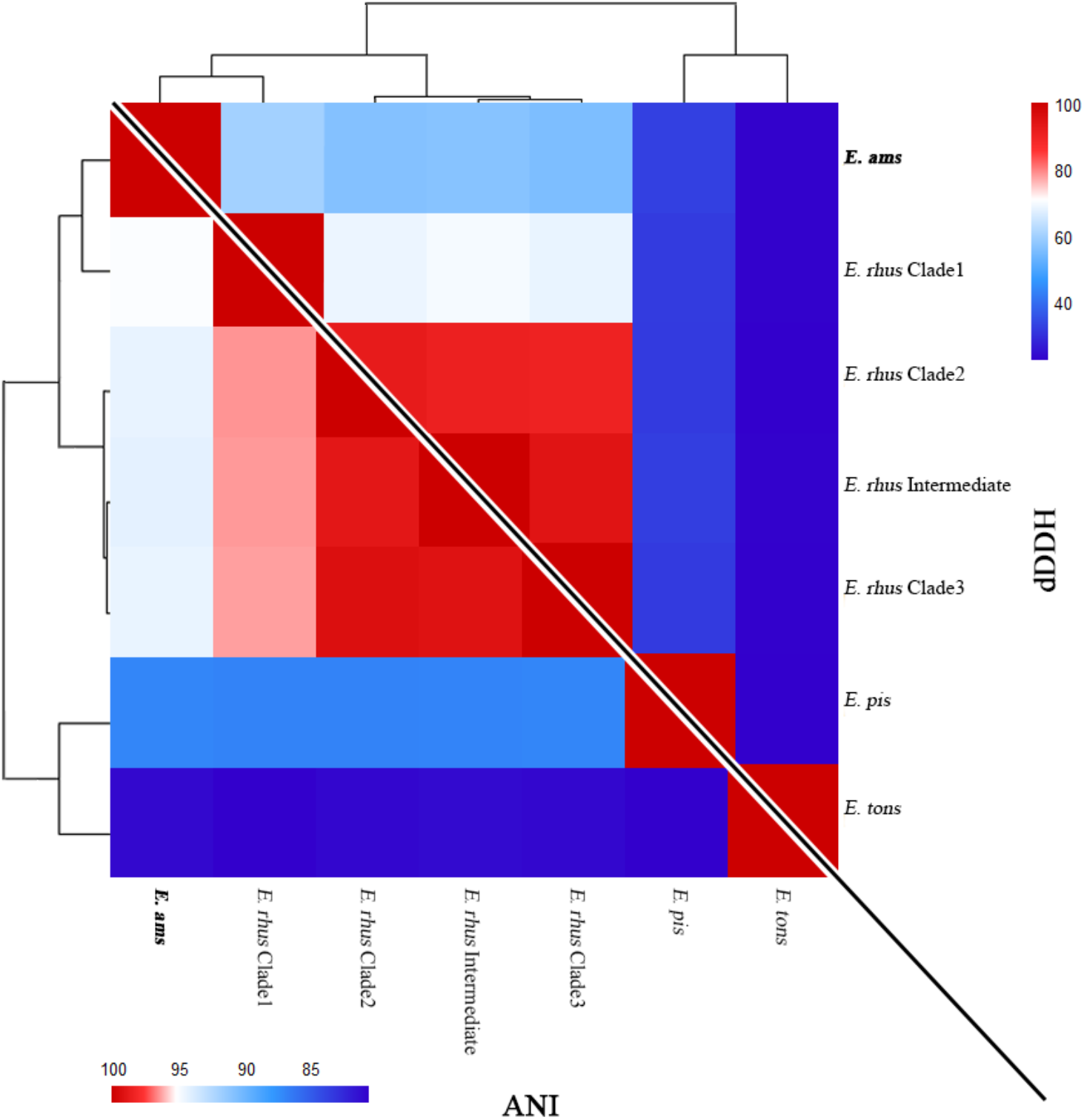
Genome relatedness of *Erysipelothrix* spp. from albatross to previously-described *Erysipelothrix* species. (A) The ANI scores comparing the *Erysipelothrix* spp. from albatross (represented by A18Y020d^T^) against other *Erysipelothrix* spp. were all below the threshold for being considered a common species (>95%). (B) The dDDH scores comparing the *Erysipelothrix* spp. from albatross against other *Erysipelothrix* spp. were also well below the accepted threshold for being considered a single species (>70%). Genomes included for comparison: *E. rhusiopathiae* Clade 1 (P-92), Clade 2 (P-190), intermediate (Fujisawa), and Clade 3 (F793); *E. piscisicarius* (15TAL0474^T^); and *E. tonsillarum* (DSM14972^T^).

Two hundred and two genes unique to the albatross-derived *Erysipelothrix* genomes (present in all 16 vs. absent in all 90 other genomes) were found using Scoary. However, after conducting blastn searches, the majority of these genes had significant hits to genes in *E. rhusiopathiae* or bacteria within the family *Erysipelotrichaceae*, or more rarely, with other bacteria. Only six genes were found with no hits or hits with low query cover (≤ 12%); a further six genes had <75% nucleotide identity with the highest scoring blastn hit (Table S3).

The *Erysipelothrix* spp. DNA from all albatross isolates produced typical amplification curves using the probe-based qPCR by Pal et al. [19] designed to detect *E. rhusiopathiae*. This is despite the fact that the forward primer and probe sequences had two mismatches each (Figure S7); based on *in silico* alignment, the 165 bp segment being amplified corresponded to positions 1,211,721 to 1,221,885 in the A18Y020d^T^ genome. The primers designed by Makino et al. [47] for *Erysipelothrix* species more broadly were a perfect match, and based on *in silico* alignment would be expected to amplify the targeted 407 bp segment of the 16S rRNA gene (positions 1,221,445 to 1,221,851 in the A18Y020d genome).

## Discussion

Through genomic comparisons, we have determined that the *Erysipelothrix* strains isolated from Indian yellow-nosed albatross chick carcasses on Amsterdam Island belong to a previously undescribed species, which we propose be named *Erysipelothrix amsterdamensis* sp. nov. This reflects its geographic origin, as well as the co-occurrence of the iconic and endemic Amsterdam albatross, *D. amsterdamensis*. Anthropogenic sources are often suspected when novel pathogens are detected in new locations. While livestock was originally suggested as a possible source of the *Erysipelothrix* spp. found on Amsterdam Island [5], we were unable to find such a link, since this particular *Erysipelothrix* species has not been previously documented elsewhere. While no earlier *Erysipelothrix* spp. isolates from this island were sequenced, using traditional phenotypic methods, initial investigations found that they belonged to serotype 1b [5]. Our finding of the same serotype using *in silico* approaches may suggest that the same species/strain has been in circulation since at least the mid-1990s. Previous PCR amplification of *Erysipelothrix* spp. isolates from Amsterdam Island was performed using the primers by Makino et al. [47] targeting the 16S rRNA gene and which should amplify sequence from all members of the genus *Erysipelothrix*. We also found that the primers and probe described by Pal et al. [19] targeting a noncoding region 3’ to the 5S rRNA gene for detection of *E. rhusiopathiae* were able to amplify DNA from *E. amsterdamensis*; care should therefore be taken when interpreting results using this primer/probe set on *Erysipelothrix* spp. isolated from rare sources. Development of a PCR protocol that distinguishes between these two species would be valuable. The genes found to be unique to *E. amsterdamensis* in this study (Table S3) would be good initial candidate targets. Moreover, the two closed genomes generated during this study will facilitate further comparative genomic studies within the genus *Erysipelothrix*. Our comparisons of the 16S rRNA and *rpoB* gene sequences from different *Erysipelothrix* species highlight the limited nucleotide diversity in the 16S rRNA gene, and confirm previous suggestions that the *rpoB* gene is a more suitable marker for *Erysipelothrix* species delineation [37].

SNP analysis of 16 *E. amsterdamensis* isolates from the three albatross chicks showed that they are highly related. Given that the number of SNPs distinguishing isolates of the same carcass was similar to that among isolates from different carcasses – as well as the close proximity in space and time of the mortality events – this strongly suggests that the three chicks were infected from a common source. Whether this small number of SNPs arose during infection within the host, or was already present in the environment, remains unknown; a molecular clock for *Erysipelothrix* has yet to be established [36]. While we cannot rule out the possibility that mutations occurred while the strains were maintained in conservation medium (several months unfrozen), we feel it is unlikely that sufficient replication occurred to explain the number of variants observed within and among these isolates. Further sampling of the albatross colony would help to determine whether this *Erysipelothrix* species was a new introduction to Amsterdam Island, and whether other strains are in circulation. Moreover, sampling from islands at comparable latitudes would help to elucidate its geographic distribution.

Interestingly, *E. amsterdamensis* harbours a *spaA* gene, which codes for what is commonly described as one of the most critical proteins for immunogenicity and virulence of *E. rhusiopathiae* [55–57]. This gene has been found to occur in all Clade 2, Clade 3 and intermediate clade *E. rhusiopathiae* genomes described to date [51], whereas *E. rhusiopathiae* Clade 1 carries a *spaB* gene [36], and *E. piscisicarius* carries a *spaC* gene [39]. That *E. amsterdamensis* was isolated in association with chick mortalities in the absence of other bacteria suggests it was the likely cause of death, and the presence of the *spaA* gene lends further support to the likely virulence of this species. However, further investigation, including challenge studies, will be necessary for such characterisation.

*E. rhusiopathiae* is typically considered an opportunistic pathogen, often manifesting clinically in stressed individuals or populations, for example following the kakapo translocation [15]. Further studies should explore whether *Erysipelothrix-*associated mortalities detected in other subantarctic islands could have been caused by the new species described here, or other novel taxa, and which set of potential host species of those relatively simple island communities may be involved in the epidemiological dynamics.

### Description of *Erysipelothrix amsterdamensis* sp. nov

*Erysipelothrix amsterdamensis* sp. nov. (am.ster.dam.en’sis. N.L. fem. *amsterdamensis* referring to Amsterdam island, from which the first strains to be characterised were isolated).

Cells are Gram-positive, rod-shaped, and non-motile. Small pin-point sized, round, flat grey colonies with smooth contours are observed on 5% horse blood agar after 24 h growth at 37°C at normal atmospheric conditions. Catalase negative. The G+C content of the genomic DNA of the type strain is 36.6%. Whole genome sequencing indicates isolates carry the *Erysipelothrix spaA* gene and belong to *Erysipelothrix* serotype 1b. Members of the species can be distinguished from other members of the genus *Erysipelothrix* based on phylogenomic analysis and genomic metrics.

The type strain, A18Y020d^T^, was isolated from an Indian yellow-nosed albatross chick carcass on Amsterdam Island. The whole genome of the type strain A18Y020d is available in GenBank (GCA_940143175).

## Author Statements

### Author contributions

Formal analysis: JZ. Investigation: JT, AC, HG, AG, TLF, TB. Methodology (Field): TB, AC, JT, AG; (Laboratory): HG, TB, AC, JT, AG; (Bioinformatics): JZ, MM, TLF. Supervision: MM, TLF, TB. Visualisation: JZ, MM. Writing – original draft: JZ, TLF. Writing – review & editing: all authors.

### Ethical approval

The experimental design was approved by the French Regional Animal Experimentation Ethical Committee n°036 (Ministry of Research permit #10257-2018011712301381) and by the Comité de l’Environnement Polaire (A-2018-123 and A 2018-139 for 2018-2019).

## Acknowledgements

We thank Nicolas Keck, Karine Lemberger, Christophe Barbraud, Karine Delord and Iain Sutcliffe for discussions related to the work. Support is acknowledged from Réserve Naturelle Nationale des Terres Australes Françaises, notably from Célia Lesage. We thank technical support agents at Ceva Biovac for help with laboratory work. This paper is a contribution of IPEV project ECOPATH-1151 to the Plan National d’Action Albatros d’Amsterdam.

## References

1. Heard MJ, Smith KF, Ripp K, Berger M, Chen J, et al. Increased threat of disease as species move towards extinction. Conserv Biol 2013;27:1378–1388.

2. Grimaldi W, Jabour J, Woehler EJ. Considerations for minimising the spread of infectious disease in Antarctic seabirds and seals. Polar Record 2011;47:56–66.

3. Cerdà-Cuéllar M, Moré E, Ayats T, Aguilera M, Muñoz-González S, et al. Do humans spread zoonotic enteric bacteria in Antarctica? Sci Total Environ 2019;654:190–196.

4. Uhart MM, Gallo L, Quintana F. Review of diseases (pathogen isolation, direct recovery and antibodies) in albatrosses and large petrels worldwide. Bird Conservation International 2018;28:169–196.

5. Weimerskirch H. Diseases threaten Southern Ocean albatrosses. Polar Biol 2004;27:374– 379.

6. Jaeger A, Lebarbenchon C, Bourret V, Bastien M, Lagadec E, et al. Avian cholera outbreaks threaten seabird species on Amsterdam Island. PLOS ONE 2018;13:e0197291.

7. Jaeger A, Gamble A, Lagadec E, Lebarbenchon C, Bourret V, et al. Impact of Annual Bacterial Epizootics on Albatross Population on a Remote Island. EcoHealth 2020;17:194– 202.

8. Bourret V, Gamble A, Tornos J, Jaeger A, Delord K, et al. Vaccination protects endangered albatross chicks against avian cholera. Conservation Letters 2018;11:e12443.

9. Gamble A, Garnier R, Jaeger A, Gantelet H, Thibault E, et al. Exposure of breeding albatrosses to the agent of avian cholera: dynamics of antibody levels and ecological implications. Oecologia 2019;189:939–949.

10. Gamble A, Bazire R, Delord K, Barbraud C, Jaeger A, et al. Predator and scavenger movements among and within endangered seabird colonies: Opportunities for pathogen spread. Journal of Applied Ecology 2020;57:367–378.

11. Opriessnig T, Coutinho TA. Erysipelas. In: Diseases of Swine. John Wiley & Sons, Ltd. pp. 835–843.

12. Bricker JM, Saif YM. Erysipelas. In: Swayne De (editor). Diseases of Poultry. Somerset, NJ: John Wiley & Sons; 2013. pp. 986–993.

13. Brooke CJ, Riley TV. Erysipelothrix rhusiopathiae: bacteriology, epidemiology and clinical manifestations of an occupational pathogen. J Med Microbiol 1999;48:789–799.

14. Kutz S, Bollinger T, Branigan M, Checkley S, Davison T, et al. Erysipelothrix rhusiopathiae associated with recent widespread muskox mortalities in the Canadian Arctic. Can Vet J 2015;56:560–563.

15. Gartrell BD, Alley MR, Mack H, Donald J, McInnes K, et al. Erysipelas in the critically endangered kakapo (Strigops habroptilus). Avian Pathol 2005;34:383–387.

16. Jayasinghe M, Midwinter A, Roe W, Vallee E, Bolwell C, et al. Seabirds as possible reservoirs of Erysipelothrix rhusiopathiae on islands used for conservation translocations in New Zealand. J Wildl Dis 2021;57:534–542.

17. Wolcott MJ. Erysipelas. In: Thomas NJ, Hunter DB, Atkinson CT (editors). Infectious Diseases of Wild Birds. Blackwell Publishing Professional. pp. 332–340.

18. Jensen WI, Cotter SE. An outbreak of erysipelas in eared grebes (Podiceps nigricollis). J Wildl Dis 1976;12:583–586.

19. Pal N, Bender JS, Opriessnig T. Rapid detection and differentiation of Erysipelothrix spp. by a novel multiplex real-time PCR assay. J Appl Microbiol 2010;108:1083–1093.

20. Bolger AM, Lohse M, Usadel B. Trimmomatic: a flexible trimmer for Illumina sequence data. Bioinformatics 2014;30:2114–2120.

21. Bankevich A, Nurk S, Antipov D, Gurevich AA, Dvorkin M, et al. SPAdes: a new genome assembly algorithm and its applications to single-cell sequencing. J Comput Biol 2012;19:455–477.

22. Gurevich A, Saveliev V, Vyahhi N, Tesler G. QUAST: quality assessment tool for genome assemblies. Bioinformatics 2013;29:1072–1075.

23. Connor TR, Loman NJ, Thompson S, Smith A, Southgate J, et al. CLIMB (the Cloud Infrastructure for Microbial Bioinformatics): an online resource for the medical microbiology community. Microbial Genomics, 2016;2:e000086.

24. Wick RR, Judd LM, Gorrie CL, Holt KE. Unicycler: Resolving bacterial genome assemblies from short and long sequencing reads. PLOS Computational Biology 2017;13:e1005595.

25. Koren S, Walenz BP, Berlin K, Miller JR, Bergman NH, et al. Canu: scalable and accurate long-read assembly via adaptive k-mer weighting and repeat separation. Genome Res 2017;27:722–736.

26. Walker BJ, Abeel T, Shea T, Priest M, Abouelliel A, et al. Pilon: an integrated tool for comprehensive microbial variant detection and genome assembly improvement. PLoS ONE 2014;9:e112963.

27. Milne I, Stephen G, Bayer M, Cock PJA, Pritchard L, et al. Using Tablet for visual exploration of second-generation sequencing data. Brief Bioinform 2013;14:193–202.

28. Seemann T. Prokka: rapid prokaryotic genome annotation. Bioinformatics 2014;30:2068–2069.

29. Conesa A, Götz S, García-Gómez JM, Terol J, Talón M, et al. Blast2GO: a universal tool for annotation, visualization and analysis in functional genomics research. Bioinformatics 2005;21:3674–3676.

30. Arndt D, Grant JR, Marcu A, Sajed T, Pon A, et al. PHASTER: a better, faster version of the PHAST phage search tool. Nucleic Acids Res 2016;44:W16–W21.

31. Carattoli A, Zankari E, García-Fernández A, Voldby Larsen M, Lund O, et al. In Silico Detection and Typing of Plasmids using PlasmidFinder and Plasmid Multilocus Sequence Typing. Antimicrobial Agents and Chemotherapy 2014;58:3895–3903.

32. Li H. Aligning sequence reads, clone sequences and assembly contigs with BWA-MEM. Oxford University Press; arXiv:1303.3997.

33. Li H, Handsaker B, Wysoker A, Fennell T, Ruan J, et al. The Sequence Alignment/Map format and SAMtools. Bioinformatics 2009;25:2078–2079.

34. Koboldt DC, Zhang Q, Larson DE, Shen D, McLellan MD, et al. VarScan 2: somatic mutation and copy number alteration discovery in cancer by exome sequencing. Genome Res 2012;22:568–576.

35. Page AJ, Cummins CA, Hunt M, Wong VK, Reuter S, et al. Roary: rapid large-scale prokaryote pan genome analysis. Bioinformatics 2015;31:3691–3693.

36. Forde T, Biek R, Zadoks R, Workentine ML, De Buck J, et al. Genomic analysis of the multi-host pathogen Erysipelothrix rhusiopathiae reveals extensive recombination as well as the existence of three generalist clades with wide geographic distribution. BMC Genomics 2016;17:461.

37. Grazziotin AL, Vidal NM, Hoepers PG, Reis TFM, Mesa D, et al. Comparative genomics of a novel clade shed light on the evolution of the genus Erysipelothrix and characterise an emerging species. Sci Rep 2021;11:3383.

38. Grazziotin AL, Vidal NM, Hoepers PG, Reis TFM, Mesa D, et al. Author Correction: Comparative genomics of a novel clade shed light on the evolution of the genus Erysipelothrix and characterise an emerging species. Sci Rep 2021;11:9861.

39. Pomaranski EK, Griffin MJ, Camus AC, Armwood AR, Shelley J, et al. Description of Erysipelothrix piscisicarius sp. nov., an emergent fish pathogen, and assessment of virulence using a tiger barb (Puntigrus tetrazona) infection model. Int J Syst Evol Microbiol 2020;70:857–867.

40. Stamatakis A. RAxML version 8: a tool for phylogenetic analysis and post-analysis of large phylogenies. Bioinformatics 2014;30:1312–1313.

41. Nylander JAA. MrModeltest. Uppsala University: Evolutionary Biology Centre; 2004.

42. Katoh K, Standley DM. MAFFT multiple sequence alignment software version 7: improvements in performance and usability. Mol Biol Evol 2013;30:772–780.

43. Yoon S-H, Ha S-M, Lim J, Kwon S, Chun J. A large-scale evaluation of algorithms to calculate average nucleotide identity. Antonie Van Leeuwenhoek 2017;110:1281–1286.

44. Meier-Kolthoff JP, Carbasse JS, Peinado-Olarte RL, Göker M. TYGS and LPSN: a database tandem for fast and reliable genome-based classification and nomenclature of prokaryotes. Nucleic Acids Research 2021;gkab902.

45. Meier-Kolthoff JP, Auch AF, Klenk H-P, Göker M. Genome sequence-based species delimitation with confidence intervals and improved distance functions. BMC Bioinformatics 2013;14:1–14.

46. Brynildsrud O, Bohlin J, Scheffer L, Eldholm V. Rapid scoring of genes in microbial pan-genome-wide association studies with Scoary. Genome Biol 2016;17:238.

47. Makino S, Okada Y, Maruyama T, Ishikawa K, Takahashi T, et al. Direct and rapid detection of Erysipelothrix rhusiopathiae DNA in animals by PCR. J Clin Microbiol 1994;32:1526–1531.

48. Kearse M, Moir R, Wilson A, Stones-Havas S, Cheung M, et al. Geneious Basic: an integrated and extendable desktop software platform for the organization and analysis of sequence data. Bioinformatics 2012;28:1647–1649.

49. Shiraiwa K, Ogawa Y, Nishikawa S, Eguchi M, Shimoji Y. Identification of serovar 1a, 1b, 2, and 5 strains of Erysipelothrix rhusiopathiae by a conventional gel-based PCR. Vet Microbiol 2018;225:101–104.

50. To H, Nagai S. Genetic and antigenic diversity of the surface protective antigen proteins of Erysipelothrix rhusiopathiae. Clin Vaccine Immunol 2007;14:813–820.

51. Forde TL, Kollanandi Ratheesh N, Harvey WT, Thomson JR, Williamson S, et al. Genomic and Immunogenic Protein Diversity of Erysipelothrix rhusiopathiae Isolated From Pigs in Great Britain: Implications for Vaccine Protection. Front Microbiol 2020;11:418.

52. Letunic I, Bork P. Interactive Tree Of Life (iTOL) v5: an online tool for phylogenetic tree display and annotation. Nucleic Acids Res 2021;49:W293–W296.

53. Jain C, Rodriguez-R LM, Phillippy AM, Konstantinidis KT, Aluru S. High throughput ANI analysis of 90K prokaryotic genomes reveals clear species boundaries. Nat Commun 2018;9:5114.

54. Kim M, Oh H-S, Park S-C, Chun J 2014. Towards a taxonomic coherence between average nucleotide identity and 16S rRNA gene sequence similarity for species demarcation of prokaryotes. International Journal of Systematic and Evolutionary Microbiology;64:346–351.

55. Makino S, Yamamoto K, Murakami S, Shirahata T, Uemura K, et al. Properties of repeat domain found in a novel protective antigen, SpaA, of Erysipelothrix rhusiopathiae. Microb Pathog 1998;25:101–109.

56. Imada Y, Goji N, Ishikawa H, Kishima M, Sekizaki T. Truncated surface protective antigen (SpaA) of Erysipelothrix rhusiopathiae serotype 1a elicits protection against challenge with serotypes 1a and 2b in pigs. Infect Immun 1999;67:4376–4382.

57. Shimoji Y, Mori Y, Fischetti VA. Immunological characterization of a protective antigen of Erysipelothrix rhusiopathiae: identification of the region responsible for protective immunity. Infect Immun 1999;67:1646–1651.

