## Supplementary material for "Genomic characterisation of a novel species of *Erysipelothrix* associated with mortalities among endangered seabirds"

**Appendix S1. Field monitoring of yellow-nosed albatross population and bacterial isolation.** As part of recent eco-epidemiological investigations implemented on Amsterdam Island in the context of repeated die-offs of yellow-nosed albatross nestlings [1–4], regular visits have been conducted to track the fate of chicks in a part of the colony where approximately 500 pairs nest (Entrecasteaux cliff, 1151 study colony; coordinates: -37.854656, 77.523990). In 2018-2019, daily visits were conducted for a month after chicks hatched (i.e., from the beginning of December) and over a series of days after chicks were one month old. As in previous years [4], a low survival of chicks (<15%) was observed (Figure S1) and an active search of dead chicks in their nests allowed necropsies to be conducted in the field before scavenging by brown rats (*Rattus norvegicus*) or brown skuas (*Stercorarius antarcticus*) occurred [3]. Sampling of the inside of organs was conducted using sterile flocked swabs stored in 1mL of liquid Amies medium in a plastic, screw cap tube (ESwab, Copan Diagnostics) until back at the field camp a few hours later (coordinates: -37.856215, 77.522724). Samples were then inoculated on Columbia nalidixic acid agar medium and stored at room temperature (circa 10-25°C). Cultures were checked daily for bacterial growth. When detected, samples of bacterial colonies from each organ were collected and stored in stock culture agar (BioRad) at room temperature until typed by MALDI-TOF several months after collection. Most colonies were eventually identified as the agent of avian cholera, *Pasteurella multocida*, based on morphology and MALDI-TOF, but the bacterial colonies from the organs of three chicks were identified as *Erysipelothrix* spp. It is those isolates that were used for sequencing in the current study.

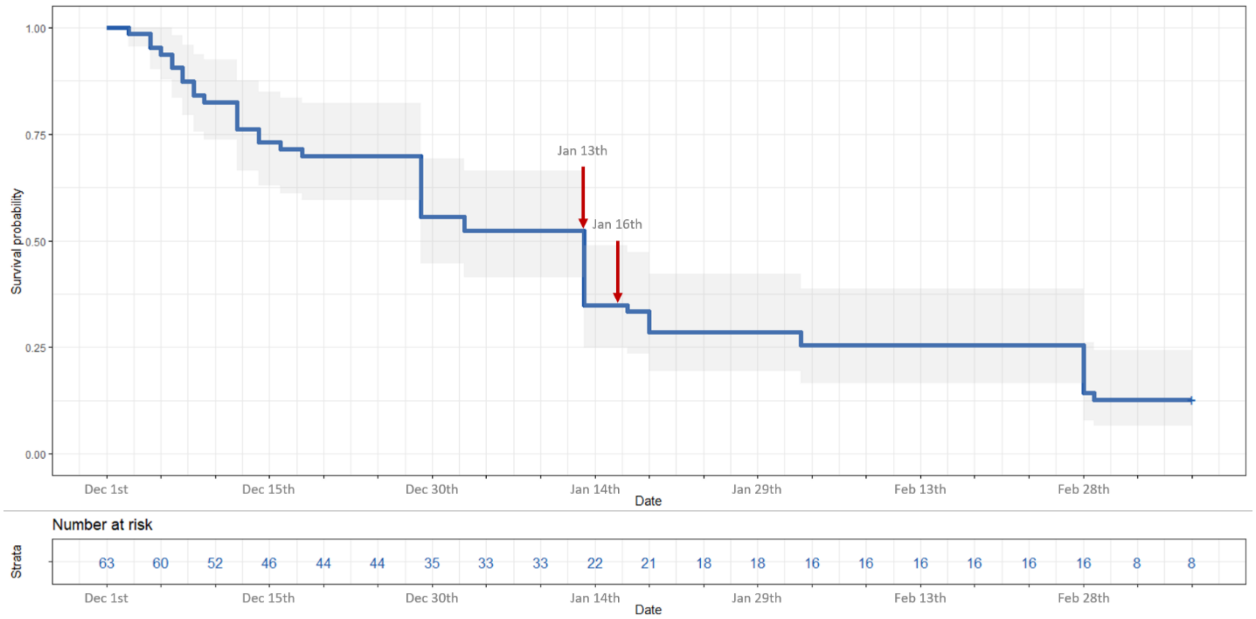


**Figure S1. Survival curve for Indian yellow-nosed albatross chicks on Amsterdam Island.** Top panel: the X-axis represents the date, from December 1st, 2018 to March 10th, 2019. The Y-axis is the survival probability. Bottom panel, in blue: number of chicks alive, displayed at a 5-day interval. The 63 chicks that were monitored were part of a control group in a vaccination experiment against *Pasteurella multocida* (causative agent of avian cholera) and therefore did not receive any treatment. Eight of those chicks were still alive and ringed on March 10th. Red arrows indicate the dates on which three chicks necropsied showed pure cultures of *Erysipelothrix* spp. from several internal organs. These chicks were found opportunistically in the monitored colony but were not part of the monitored group.

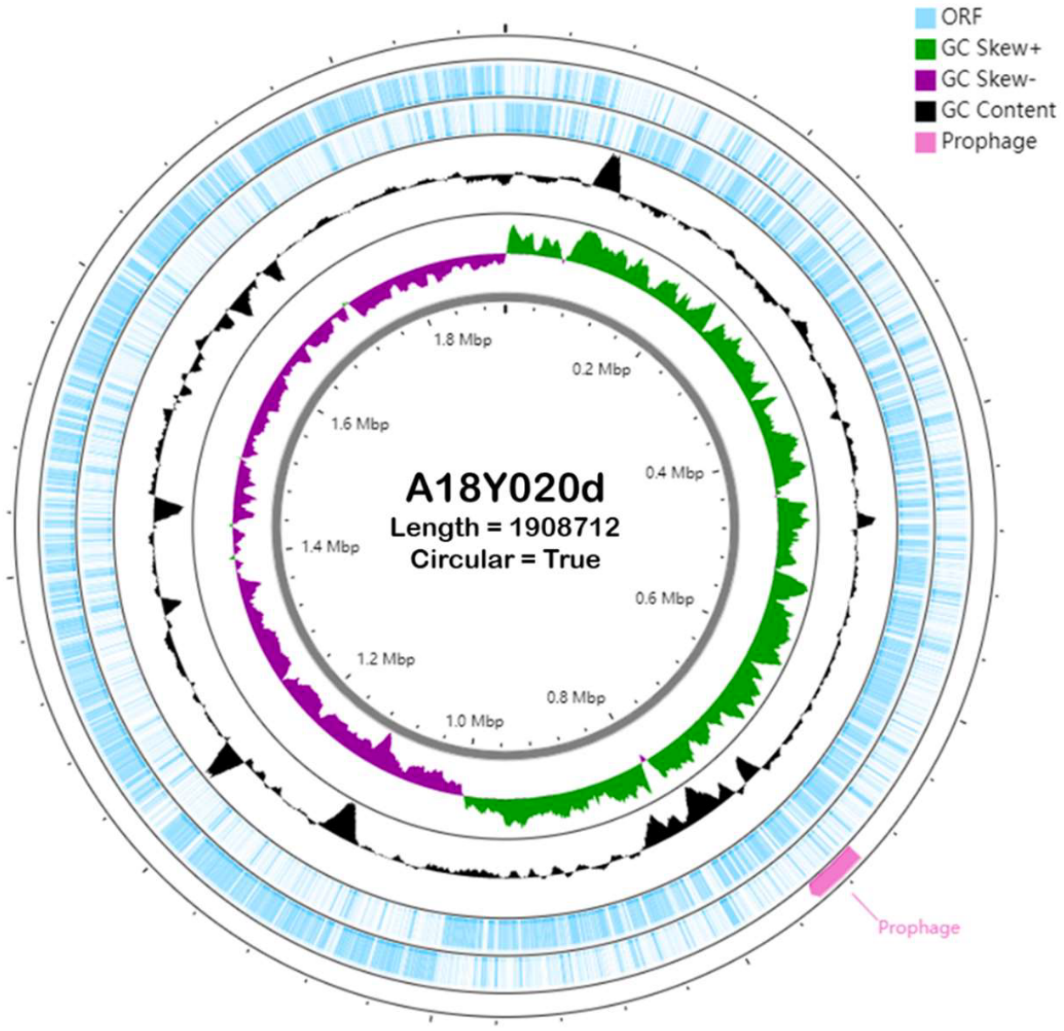


**Figure S2. Circular representation of *E. amsterdamensis* sp. nov. type strain A18Y020d$^{T}$ genome.** Circles from inside to outside represent (i) nucleotide position of genome (grey), (ii) GC Skew+ region with G content greater than C (green) and GC skew- region with G content less than C (purple), (iii) G + C content (black), (iv) predicted open reading frames (ORFs) (blue, negative strand – inner, positive strand – outer), and (v) prophage annotated with PHASTER (pink), respectively. The figure was plotted using CGView server (http://cgview.ca/) [5].

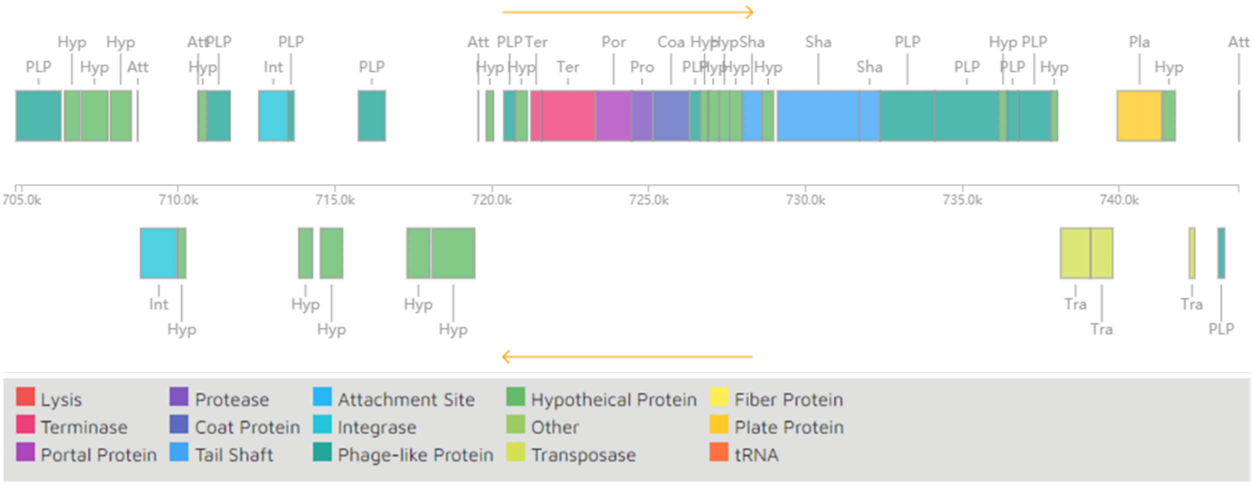


**Figure S3**. *Erysipelothrix amsterdamensis* sp. nov. strain A18Y020d$^{T}$ prophage schematic diagram plotted by PHASTER. The position of prophage in A18Y020d$^{T}$ genome ranged from base pairs 704,829 to 743,808. The upper and lower parts in the figure represent forward and reverse genes, respectively.

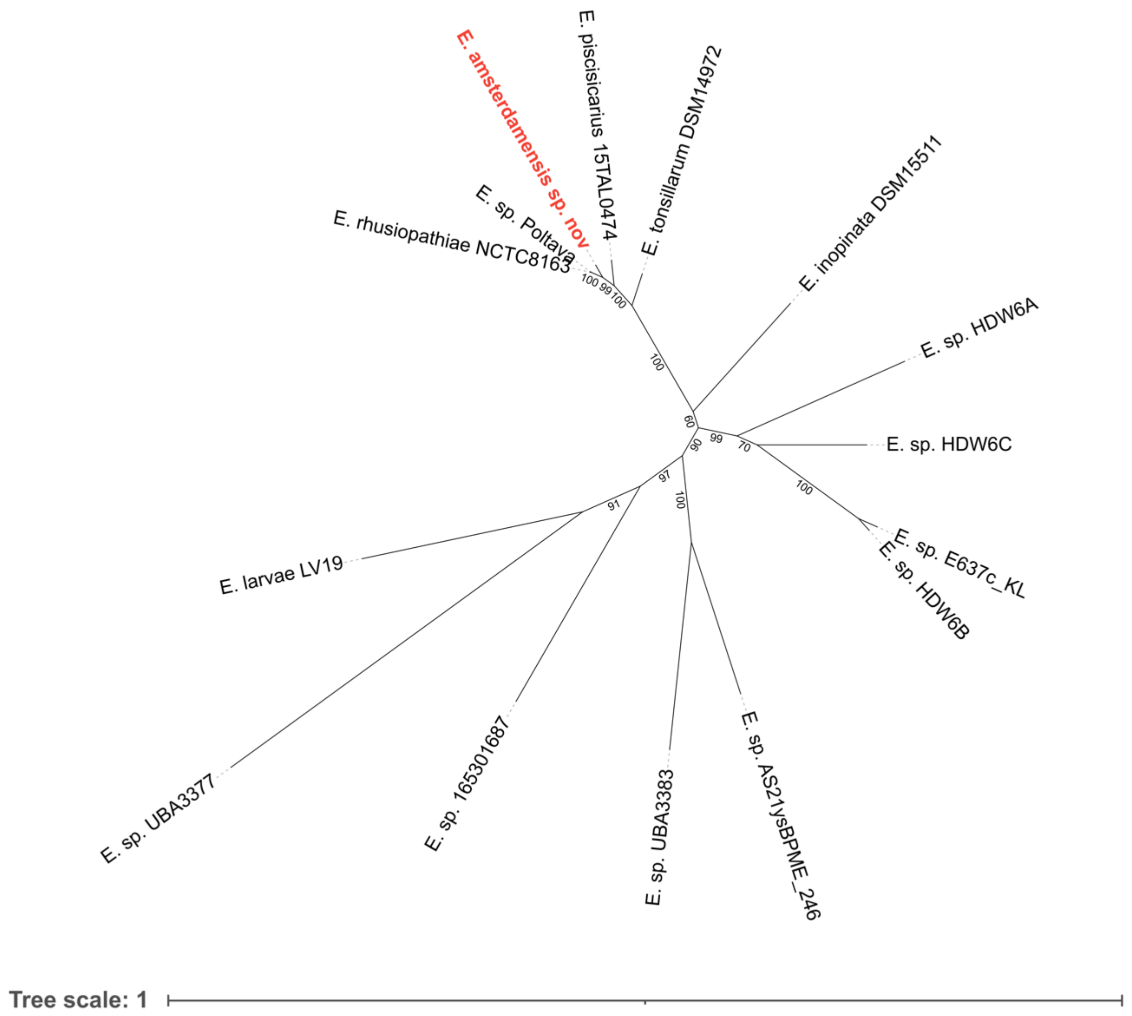


**Figure S4. Phylogenetic tree of *Erysipelothrix* species based on *rpoB* gene sequences.** This confirms that *E. rhusiopathiae, E. piscisicarius* and *E. tonsillarum* are the most closely related species to the *Erysipelothrix* spp. isolated from albatrosses in this study. Nine other yet-to-be-named species deposited in the Genome Taxonomy Database (https://gtdb.ecogenomic.org/), along with *E. inopinata* and *E. larvae*, are more distantly related. *E.* sp. Poltava appears to be a strain of *E. rhusiopathiae* based on further analyses (Figure S5 and S6).

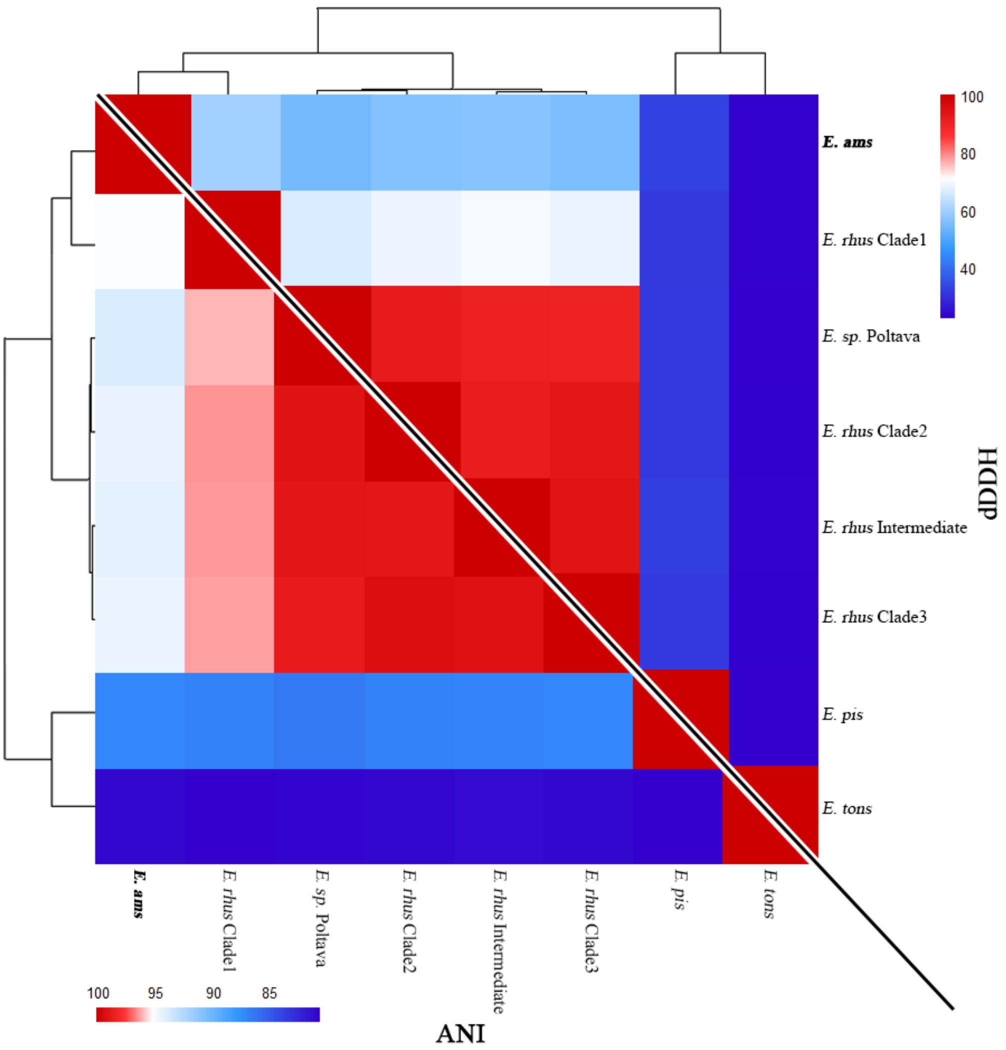


**Figure S5. Genome relatedness among *Erysipelothrix* species.** As shown in Figure 3, but with inclusion of recently deposited *Erysipelothrix* sp. Poltava (GCA_023221615.1) for comparison. Based on accepted thresholds for ANI (>95%) and dDDH scores (>70%) for inclusion in the same species, *Erysipelothrix* sp. Poltava would be classified as *Erysipelothrix rhusiopathiae.*

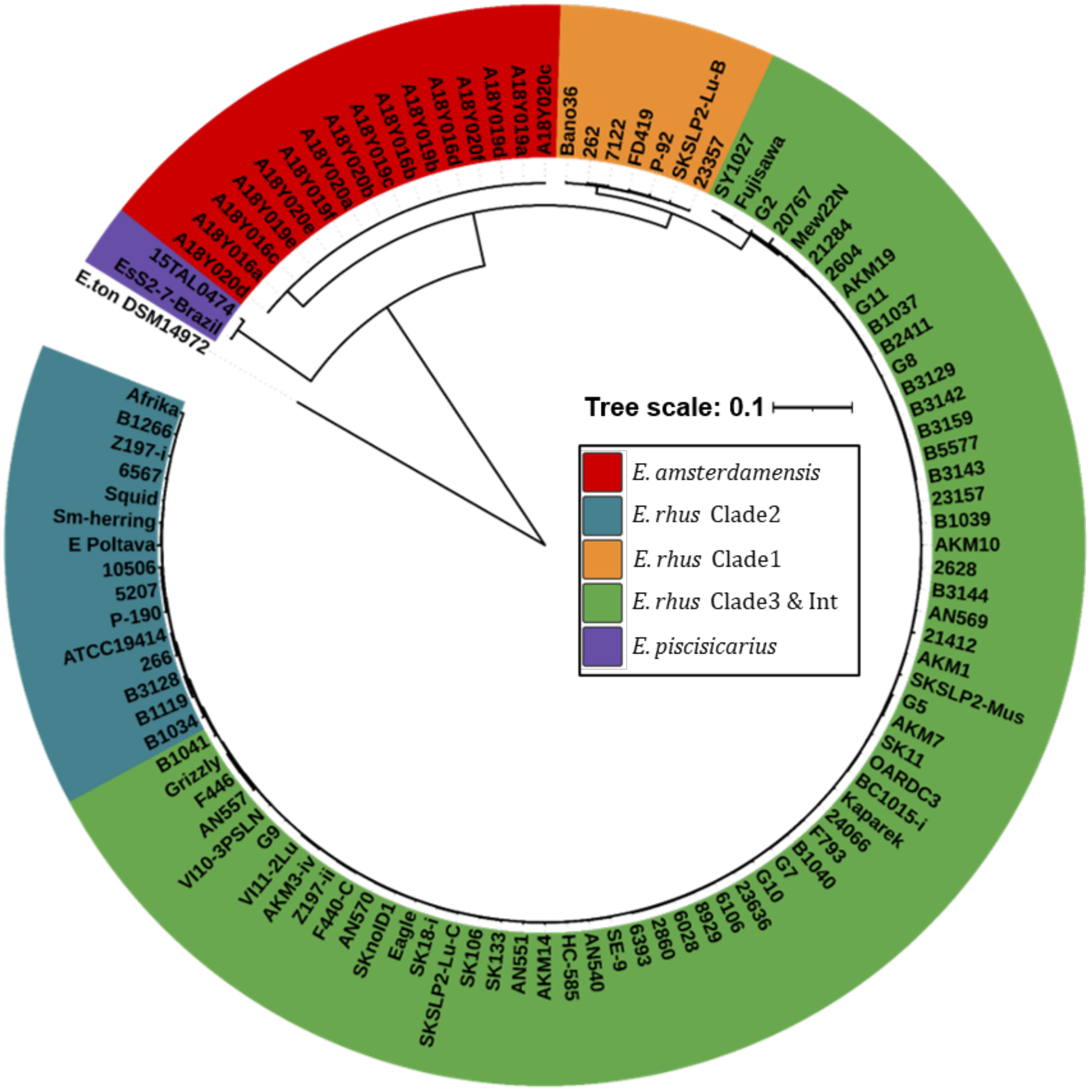


**Figure S6. Phylogenomic comparison of *Erysipelothrix* genomes.** Core genome phylogeny rooted to *E. tonsillarum* strain DSM14972$^{T}$. Genomes of *E. piscisicarius* (purple), *E. amsterdamensis* sp. nov (this study; pink), and *E. rhusiopathiae* Clade 1 (yellow), Clade 2 (blue) and Clade 3/Intermediate (green) are shown. The recently deposited *Erysipelothrix* sp. Poltava (GCA_023221615.1) clusters with genomes of *E. rhusiopathiae* Clade 2.

5’-ATTTCTCTAGCAGGTGATTTGG-3’ – Pal *et al*. (2010) Ery4423F (forward primer)
ATTTCCCTAGCGGGTGATTTGG – sequence in albatross isolates

5’-FAM-AACGAAACGATTAGTAGTCCAACA-BHQ-3’ – Pal *et al*. (2010) *E. rhusiopathiae* probe
GACGAAACGATTAGTAGTCCGACA – sequence in albatross isolates

**Figure S7. Nucleotide discrepancies between published primer and probe sequences for *E. rhusiopathiae* and sequences found within the *E. amsterdamensis* sp. nov. genome.** Primer/probe sequences are based on those by Pal et al., 2010 [6].

**Table S1. *Erysipelothrix* spp. isolates recovered and sequenced from three Indian yellow-nosed albatross chick carcasses on Amsterdam Island.** Number of reads and mean depth of coverage are based on paired-end Illumina data. Number of single nucleotide polymorphisms (SNPs) is based on mapping to the closed genome of strain A18Y020d$^T$ (*) obtained through hybrid assembly. The SNPs identified in this latter isolate are in comparison to the other 15 genomes analysed.

| carcass ID | date sampled (dd-mm-yyyy) | isolate ID | organ | # reads | mean coverage (X) | # MinION reads | # SNPs | Accession |
|---|---|---|---|---|---|---|---|---|
| AMS18-NEC-016 | 13-01-2019 | A18Y016a | liver | 359262 | 95 | 21074 | 3 | ERS11482829 |
| | | A18Y016b | lung | 434748 | 112 | NA | 3 | ERS11482830 |
| | | A18Y016c | brain | 469941 | 127 | NA | 5 | ERS11482831 |
| | | A18Y016d | heart | 413222 | 108 | NA | 2 | ERS11482832 |
| AMS18-NEC-019 | 16-01-2019 | A18Y019a | liver | 388054 | 101 | NA | 2 | ERS11482833 |
| | | A18Y019b | heart | 582025 | 148 | NA | 8 | ERS11482834 |
| | | A18Y019c | brain | 387940 | 101 | NA | 5 | ERS11482835 |
| | | A18Y019d | liver | 219988 | 58 | NA | 2 | ERS11482836 |
| | | A18Y019e | lung | 217733 | 58 | NA | 1 | ERS11482837 |
| | | A18Y019f | brain | 337715 | 91 | NA | 4 | ERS11482838 |
| AMS18-NEC-020 | 16-01-2019 | A18Y020a | liver | 263781 | 70 | NA | 1 | ERS11482839 |
| | | A18Y020b | heart | 338940 | 90 | NA | 3 | ERS11482840 |
| | | A18Y020c | brain | 276491 | 73 | NA | 3 | ERS11482841 |
| | | A18Y020d* | liver | 394696 | 101 | 104217 | 3 | ERS11482842 |
| | | A18Y020e | heart | 315117 | 81 | NA | 5 | ERS11482843 |
| | | A18Y020f | lung | 336890 | 85 | NA | 3 | ERS11482844 |

NA: not applicable

**Table S2. Comparison of two closed *Erysipelothrix* spp. genomes from albatrosses (A18Y016a and A18Y020d$^{T}$) with the *E. rhusiopathiae* Fujisawa genome.** Values for the Fujisawa genome are shown using the same annotation software as implemented for the albatross genomes (Prokka) to facilitate direct comparison, and those published in the manuscript by Ogawa et al (2011) originally describing this genome [7].

| **Genome** | **A18Y016a** | **A18Y020d$^{+}$** | **Fujisawa (Prokka)** | **Fujisawa (Ogawa)** |
|---|---|---|---|---|
| Total Genome Size | 1910750 bp | 1908712 bp | 1787941 bp | |
| Accession number | GCA_940143155 | GCA_940143175 | | GCF_000270085.1 |
| Prophage Size | 38981 bp | 38980 bp | 41304 bp | |
| Location of prophage in genome* | 1004029 – 1043009 | 704829 – 743808 | 624715 – 666018 | 625920 – 662139 |
| GC Content | 36.52% | 36.52% | 36.56% | |
| Predicted Genes (Prokka) | 1855 | 1848 | 1755 | 1704 |
| tRNA | 55 | 49 | 55 | |
| tmRNA | 1 | 1 | 1 | |
| rRNA | 20 | 21 | 21 | |
| CDS | 1779 | 1777 | 1678 | |

* Different phage positions are the result of different start and end positions of the two circular genomes. The genome structure is identical.
+ designated as type strain

**Table S3. Genes unique to *Erysipelothrix amsterdamensis* sp. nov.** Results are based on output from Roary/Scoary, and confirmed through blastn searches. Genes shaded in grey produced best blastn hits with percent identity below 75%.

| Locus tag | Position | Gene name* / Description | Query cover | E value | Percent Identity |
|---|---|---|---|---|---|
| ERYAMS_00687 | 717290-718012 | hypothetical protein | 7% | 9e-04 | 85.71% |
| ERYAMS_00688 | 718084-719445 | hypothetical protein | 5% | 0.002 | 77.11% |
| ERYAMS_00715 | 739934-741355 | *rnhA* / viroplasmin family protein | - | - | - |
| ERYAMS_00716 | 741387-741785 | hypothetical protein | 12% | 0.002 | 85.71% |
| ERYAMS_00739 | 766929-767705 | hypothetical protein | - | - | - |
| ERYAMS_01411 | 1450615-1451079 | hypothetical protein | 7% | 0.024 | 91.89% |
| ERYAMS_00712 | 737822-738031 | hemolysin XhlA family protein | 100% | 2e-24 | 73.93% |
| ERYAMS_00745 | 783568-784743 | restriction endonuclease subunit S | 61% | 1e-119 | 73.86% |
| ERYAMS_01042 | 1103769-1105889 | *lag D* / peptidase domain-containing ABC transporter | 99% | 0.0 | 68.24% |
| ERYAMS_01043 | 1105901-1109128 | type 2 lantipeptide synthetase LanM family protein | 90% | 0.0 | 69.01% |
| ERYAMS_01410 | 1448445-1449704 | putative DNA binding domain-containing protein | 78% | 6e-85 | 67.83% |
| ERYAMS_01413 | 1453839-1454153 | 4-hydroxybenzoate polyprenyltransferase | 84% | 3e-11 | 67.91% |

*provided where relevant
Where no values shown (-), no significant similarity found.

**Supplementary References**

1. **Bourret V, Gamble A, Tornos J, Jaeger A, Delord K, *et al.*** Vaccination protects endangered albatross chicks against avian cholera. *Conservation Letters* 2018;11:e12443.

2. **Gamble A, Garnier R, Jaeger A, Gantelet H, Thibault E, *et al.*** Exposure of breeding albatrosses to the agent of avian cholera: dynamics of antibody levels and ecological implications. *Oecologia* 2019;189:939–949.

3. **Gamble A, Bazire R, Delord K, Barbraud C, Jaeger A, *et al.*** Predator and scavenger movements among and within endangered seabird colonies: Opportunities for pathogen spread. *Journal of Applied Ecology* 2020;57:367–378.

4. **Jaeger A, Gamble A, Lagadec E, Lebarbenchon C, Bourret V, *et al.*** Impact of Annual Bacterial Epizootics on Albatross Population on a Remote Island. *EcoHealth* 2020;17:194–202.

5. **Grant JR, Stothard P**. The CGView Server: a comparative genomics tool for circular genomes. *Nucleic Acids Res* 2008;36:W181-184.

6. **Pal N, Bender JS, Opriessnig T**. Rapid detection and differentiation of Erysipelothrix spp. by a novel multiplex real-time PCR assay. *J Appl Microbiol* 2010;108:1083–1093.

7. **Ogawa Y, Ooka T, Shi F, Ogura Y, Nakayama K, *et al.*** The genome of Erysipelothrix rhusiopathiae, the causative agent of swine erysipelas, reveals new insights into the evolution of firmicutes and the organism's intracellular adaptations. *J Bacteriol* 2011;193:2959–2971.
